# *Aedes aegypti* in Maryland: The need for elevated vector surveillance at the face of a dynamic climate

**DOI:** 10.1101/2023.10.02.560479

**Authors:** Roy Faiman, Autumn Goodwin, Jaykob Cave-Stevens, Alyssa Schultz, Jewell Brey, Tristan Ford

## Abstract

We report the collection of three *Aedes aegypti* adult female mosquitoes in a Rockville, Maryland backyard in late July, 2023, followed by the emergence of 15 adults collected as larvae in a residential backyard in Baltimore, Maryland in mid-September, 2023. *Aedes aegypti* is a species primarily associated with tropical and subtropical regions, known for its significance as a vector of arboviruses, including Yellow Fever, Dengue, Zika, and Chikungunya among others. In the continental United States, *Ae. aegypti* populations are mostly found in the southeast, and in several isolated locations such as southern California and Arizona. This finding raises questions about the potential establishment and survival of *Ae. aegypti* in temperate climates, emphasizing the critical importance of robust vector surveillance programs in preventing potential outbreaks of vector-borne diseases in regions not traditionally considered endemic for this species.

## Introduction

Originating in Africa, *Aedes aegypti* found its way to the Americas through inadvertent transportation on ships bound for the New World during the Transatlantic Slave Trade. This mosquito species was once abundant and firmly rooted in most of the eastern United States. However, in the 1980s, the introduction of *Ae. albopictus* in the U.S. led to the displacement of *Ae. aegypti* across much of its former range. Presently, *Ae. aegypti* is primarily confined to Southern Florida and Texas (Burkett-Cadena, 2013), with occasional established populations reported in other regions in the U.S. (Gloria-Soria et al., 2014, 2022; Lima et al., 2016; Merrill et al., 2005). *Aedes aegypti* is a medically significant mosquito species known for its association with the transmission of several arboviruses of acute global health concern (Gubler, 2011), including Yellow Fever, Dengue, Zika, and Chikungunya among others. *Aedes aegypti*’s range is contingent on temperature, as this species is unable to endure prolonged periods of cold, such as those experienced in temperate winters (Farajollahi and Crans, 2012). Christophers defined the geographical boundaries for year-round activity, specifying locations with a mean temperature of 10 °C or higher (Christophers, 1960). This encompasses latitudes between 30-35° N and S during the coldest month. This species’ presence in traditionally considered temperate zones is a concerning development, as it increases the risk of local transmission of diseases previously considered exotic to these areas (Ryan at al., 2019). Historically, this species has been linked to tropical and subtropical regions, and its presence in temperate climates is sporadic (Hahn et al., 2016; Hahn et al., 2017). This report highlights the collection and identification of *Ae. aegypti* mosquitoes in two separate urban residential backyards in Maryland, signifying the necessity of comprehensive and updated vector surveillance.

## Methods

Adult females were collected by R.F. using a mouth aspirator in a Rockville, Maryland neighborhood in Late July, 2023. Mosquito larvae were collected by A.G. from water-holding containers in the backyard of a residence located in the Hampden neighborhood of Baltimore, Maryland in mid-September, 2023. Larvae were reared until adult emergence indoors in office conditions, and adult mosquitoes were provided with 10% sucrose. Species identification was confirmed through morphological examination by an entomologist, followed by imaging on an IDX vector identification device (IDX. Vectech Inc. Baltimore, MD) (Fig. 1) (Brey, 2022).

**Figure 1.**
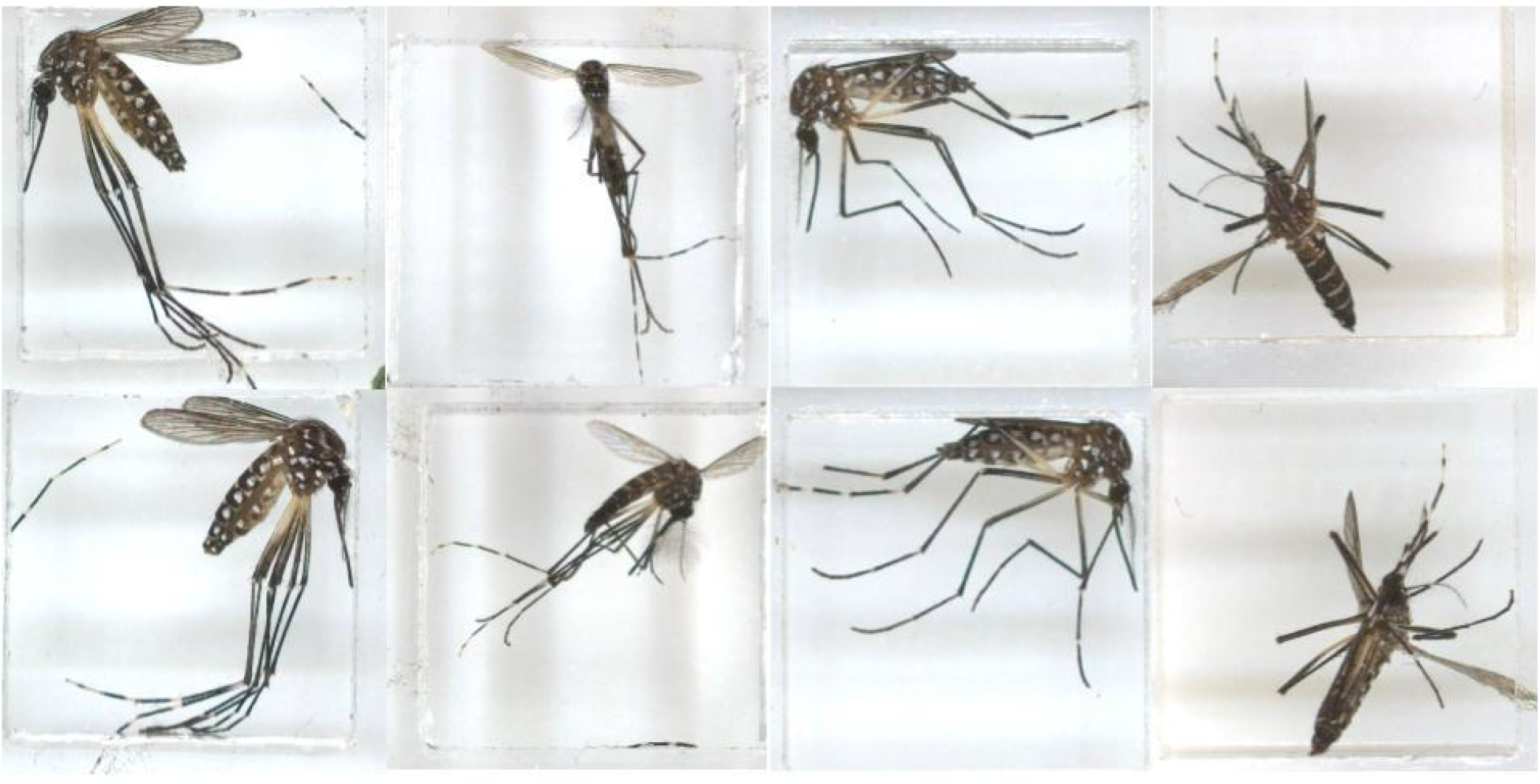
Aedes aegypti adults. Two specimens collected in Baltimore, MD (left; female and male), and two specimens collected in Rockville, MD (right; two females). Each specimen was imaged from two sides (top and bottom rows). Images were produced on an IDX device.

## Results and Discussion

In Rockville, three adult females were collected by live aspiration (Human Landing Catches) and soon were identified as *Ae. aegypti* using a stereo microscope. In Baltimore, 15 adult mosquitoes emerged from the collected larvae, and subsequent morphological examination confirmed them all to be *Ae. aegypti*. From this batch of larvae emerged 51 *Ae. albopictus* as well. *Aedes aegypti* is recognized by characteristic black and white markings on its body, as well as the distinctive white lyre-shaped pattern on the dorsal thorax (versus the single white stripe found on *Ae. albopictus*, or golden lyre and long palps on *Ae. japonicus*), as well as presenting silverish scales on the clypeus and characteristic short, rectangular white banding on the abdominal tergites (Burkett-Cadena, 2013). All adults were imaged on an IDX, further confirming their species identifications (Fig. 1) (Brey, 2022).

Although sporadically reported from locations beyond its northern distribution limits (Hahn et al., 2016; Hahn et al., 2017), the emergence of *Ae. aegypti* from collected larvae in a temperate climate like Baltimore and Rockville, Maryland, though not the first, is an uncommon finding which may be rooted in limited surveillance programs in the region. This species is predominantly associated with warmer, tropical regions found in the southeastern US, though a resident population was recently reported in Washington D.C. (Capitol Hill neighborhood), thought to be overwintering in subterranean refugia (Lima et al., 2016, Gloria-Soria et al., 2018), in the same region as the detections in this report. Both Baltimore and Rockville, Maryland similarly maintain potential habitats for overwintering, such as subterranean metro stations, larger-scale sewer mains, storm drains, and basements (Lima et al., 2016).

The repeated presence of this species in Maryland (and D.C.) raises questions about its potential establishment and survival in temperate climates, which may be influenced by factors such as climate change, urbanization, and the availability of suitable breeding and overwintering habitats. Based on these factors, predictions of future expansion of this species’ distribution suggest it will likely become established in northeastern climes over the coming years and decades (Laporta et al., 2023).

## Conclusion

The finding of *Ae. aegypti* mosquitoes in Maryland highlight the need for vigilant vector surveillance and control measures. Early detection and monitoring are crucial for preventing the establishment of arboviral diseases in new regions. This discovery serves as a stark reminder of the evolving threat posed by vector-borne diseases and underscores the importance of sustained efforts in vector surveillance and control in a dynamic climate.

## Notes

### Competing Interest Statement

The authors have declared no competing interest.

## References

Burkett-Cadena, Nathan D. “Mosquitoes of the southeastern United States.” University of Alabama Press, 2013.

Christophers, Samuel Rickard. “Aedes aegypti: the yellow fever mosquito.” CUP Archive, 1960.

Farajollahi, Ary, and Scott C. Crans. “A checklist of the mosquitoes of New Jersey with notes on established invasive species.” Journal of the American Mosquito Control Association 28.3 (2012): 237–239.

Gloria-Soria, Andrea, et al. “Origin of the dengue fever mosquito, Aedes aegypti, in California.” PLoS Neglected Tropical Diseases 8.7 (2014): e3029.

Gloria-Soria A, Lima A, Lovin DD, Cunningham JM, Severson DW, Powell JR. “Origin of a high-latitude population of Aedes aegypti in Washington, DC.” The American journal of tropical medicine and hygiene. 2018 Feb;98(2):445.

Gloria-Soria, Andrea, et al. “Origins of high latitude introductions of Aedes aegypti to Nebraska and Utah during 2019.” Infection, Genetics and Evolution 103 (2022): 105333.

Gubler Duane J. “Dengue, urbanization and globalization: the unholy trinity of the 21st century.” Tropical medicine and health 39.4SUPPLEMENT (2011): S3–S11.

Hahn, Micah B., et al. “Reported distribution of Aedes (stegomyia) aegypti and Aedes (stegomyia) albopictus in the United States, 1995-2016 (diptera: Culicidae).” Journal of medical entomology 53.5 (2016): 1169–1175.

Hahn Micah B., et al. “Updated reported distribution of Aedes (stegomyia) aegypti and Aedes (stegomyia) albopictus (diptera: Culicidae) in the United States, 1995–2016.” Journal of medical entomology 54.5 (2017): 1420–1424.

Laporta, Gabriel Z., et al. “Global distribution of Aedes aegypti and Aedes albopictus in a climate change scenario of regional rivalry.” Insects 14.1 (2023): 49.

Lima A, Lovin DD, Hickner PV, Severson DW. “Evidence for an Overwintering Population of Aedes aegypti in Capitol Hill Neighborhood, Washington, DC.” Am J Trop Med Hyg. 2016 Jan;94(1):231–5. doi: 10.4269/ajtmh.15-0351. Epub 2015 Nov 2. PMID: 26526922; PMCID: PMC4710436.

Merrill Samuel A.; Ramberg Frank B.; and Hagedorn Henry H., “Phylogeography and population structure of Aedes aegypti in Arizona.” American Journal of Tropical Medicine and Hygiene. 72, 3. 304–310. (2005). https://digitalcommons.wustl.edu/open_access_pubs/1740

Ryan Sadie J., et al. “Global expansion and redistribution of Aedes-borne virus transmission risk with climate change.” PLoS neglected tropical diseases 13.3 (2019): e0007213.

